# Interindividual differences in pain can be explained by fMRI, sociodemographic, and psychological factors

**DOI:** 10.1101/2023.07.06.547919

**Authors:** Suhwan Gim, Dong Hee Lee, Sungwoo Lee, Choong-Wan Woo

## Abstract

In a recent article, Hoeppli et al. (2022) reported that sociodemographic and psychological factors were not associated with interindividual differences in reported pain intensity. In addition, the interindividual differences in pain could not be detected by thermal pain-evoked brain activities measured by functional Magnetic Resonance Imaging (fMRI). Their comprehensive analyses provided convincing evidence for these null findings, but here we provide another look at their conclusions by analyzing their behavioral data and a large-scale fMRI dataset involving thermal pain (*N* = 124). Our main findings are as follows: First, a multiple regression model incorporating all available sociodemographic and psychological measures could significantly predict the interindividual differences in reported pain intensity. The key to achieving a significant prediction was including multiple individual difference measures in a single model. Second, with fMRI data from a relatively homogeneous group of 124 participants, we could identify brain regions and a multivariate pattern-based predictive model significantly correlated with the interindividual differences in reported pain intensity. Our results, along with the findings of Hoeppli et al., highlight the challenge of predicting interindividual differences in pain, but also suggest that it is not an impossible task.

---

Developing neuroimaging biomarkers of pain has the potential to improve pain assessment and management by providing objective measures of the subjective experience of pain^1^. However, it remains challenging to develop such biomarkers due to the substantial interindividual variability in brain systems for pain processing^2^ and pain-expressive behaviors^3^. This interindividual variability is influenced by multiple factors, including biological, psychological, and social ones, requiring systematic investigation of how these multiple factors and components are associated with interindividual variability in pain. In this endeavor, Hoeppli et al.’s recent report could be discouraging to those hoping for progress in brain-based pain biomarker development. The study provided two main conclusions, one for the sociodemographic and psychological factors and the other for the fMRI signal. They showed that both data types failed to explain interindividual differences in pain sensitivity and reported pain intensity.

First, they reported that individual differences in pain could not be explained by sociodemographic and psychological factors, which is somewhat inconsistent with what has been known about their effects on pain^4^. For example, previous studies have shown that pain experience can be influenced by age^5^, ethnicity^6^, sex^7^, and emotional states^8^, among many.Potentially, the inconsistency may come from the fact that they did not consider the complex interactions among the sociodemographic and psychological factors. Though they examined the relationship between pain ratings and each of those factors separately, it is widely recognized that sociodemographic and psychological factors are intercorrelated, and their influences on pain are likely to come from their complex interactions. For example, a previous study reported that the ethnicity effects on pain were mediated by perceived discrimination^9^, and another study showed that psychological and personality factors identified to be important for chronic pain were also associated with the socioeconomic status of patients with chronic pain^10^. These highlight that sociodemographic and psychological factors interact with each other to influence pain. Thus, to better understand the effects of sociodemographic and psychological factors on interindividual differences in pain, it is crucial to test their combined effects, for example, by incorporating them in a single model^11^.

To test this idea, we reanalyzed the behavioral data from Hoeppli et al. to examine the complex interactions between sociodemographic and psychological factors and their combined effects on interindividual differences in self-reported pain intensity. Specifically, we first examined the correlations between the sociodemographic and psychological measures and then performed a multiple regression with cross-validation using the sociodemographic and psychological measures as independent variables. The result showed that the sociodemographic and psychological measures were highly intercorrelated; 20% of all pairs (11 out of 55 pairs) showed significant correlations at *q* < 0.05, false discovery rate (FDR) corrected (**Fig. 1a**). This supports the idea that the sociodemographic and psychological variables are highly inter-connected. When we examined the combined effects of the sociodemographic and psychological measures on pain ratings using multiple regression with leave-one-subject-out cross-validation (LOSO-CV), the prediction-outcome correlation was significant, *r* = 0.260, *p* = 0.026. To account for potential issues related to multicollinearity, we also tested a principal component regression with LOSO-CV, and the results were comparable, *r* = 0.270, *p* = 0.021. Importantly, when we reduced the number of predictors to two or one, the cross-validated correlations were reduced to negative values (mean *r* = -0.005 and -0.1608 for two factors and one factor; **Fig. 1b**). Overall, different from Hoeppli et al.’s report, our findings show that the sociodemographic and psychological factors can explain individual differences in pain ratings, but only when their combined effects were accounted for.

**Fig. 1.**
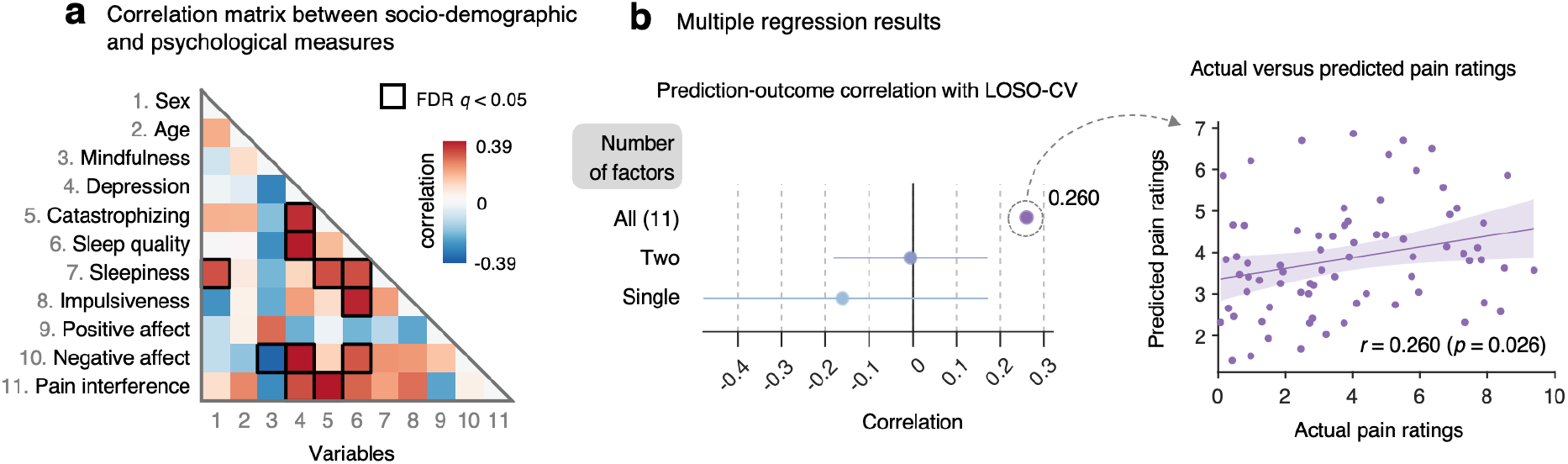
Reanalysis of behavioral data from Hoeppli et al. We reanalyzed data provided by the authors. We only included participants’ data with no missing values (*N* = 73) in this analysis. **a**, The correlations between the demographic and psychological measures showed a large number of significant correlations (11 out of 55 pairs) after correcting for multiple comparisons, *q* < 0.05, false discovery rate (FDR). **b**, Results of multiple regression analysis with leave-one-subject-out cross-validation (LOSO-CV). (Left) We examined the cross-validated prediction performance of linear regression models with varied numbers of independent variables (here, one, two, and all eleven variables). The cross-validated prediction-outcome correlations were smaller than zero for the model with a single independent variable, mean ± SD of *r* = -0.161 ± 0.334, and for the model with two independent variables, *r* = -0.005 ± 0.176. However, the multiple regression model with all 11 variables showed a significant cross-validated prediction-outcome correlation, *r* = 0.260, *p* = 0.0263. Each dot represents the mean prediction-outcome correlation, and the error bar represents its standard deviation. (right) The scatter plot shows the actual versus cross-validated predicted pain ratings based on the multiple regression model with all 11 independent variables.

Second, Hoeppli et al. reported that the individual differences in pain ratings could not be detected by fMRI signal, which is also somewhat inconsistent with what has been known about pain and the brain. For example, two recent studies have shown that multivariate whole-brain functional connectivity patterns can predict the interindividual differences in pain of patients with chronic pain^12^ or healthy participants^13^. One of the functional connectivity models also showed a significant association with the number of pain sites in a large-scale dataset in an independent study^14^. Furthermore, a previous study showed that the striatal fMRI activity could explain increased pain ratings in African Americans compared to Hispanics and non-Hispanic white^9^, which was also associated with perceived discrimination. In addition, another study reported that multivariate functional connectivity patterns related to pain-predictive psychological traits were associated with the socioeconomic status of patients with chronic pain^10^. All these findings suggest that fMRI signals can explain interindividual differences in pain to some extent.

To reconcile the discrepancy, we analyzed a large-scale pain fMRI dataset (*N* = 124) that included a similar number of participants to Hoeppli et al. We aimed to replicate their main findings that there was no brain region or multivariate pattern that could explain interindividual differences in pain ratings. We first examined the distribution of average pain intensity ratings for the high-intensity heat stimuli (47.5°C). As shown in **Fig. 2a**, the ratings showed a wide range of distribution, which was consistent with Hoeppli et al—the averaged pain ratings ranged from “Moderate” to near “Strongest imaginable” on the general Labeled Magnitude Scale (gLMS)^15^. Then, we analyzed fMRI data using the same methods that Hoeppli et al. used in their study; 1) univariate general linear model, where we included interindividual variations in pain ratings as a covariate, and 2) multivariate lasso-regularized principal component regression (LASSO-PCR) to predict interindividual variations in pain ratings. Though the analysis methods were the same, the results were quite different. The univariate analysis identified multiple brain regions (**Fig. 2b**), including periaqueductal gray (PAG) and supplementary motor area (SMA), and visual cortex, significantly correlated with interindividual variations in pain ratings at *q* < 0.05, FDR corrected. The multivariate predictive modeling with LASSO-PCR also showed a significant prediction-outcome correlation with LOSO-CV, *r* = 0.252, *p* = 0.0047 (**Fig. 2c**).Ventrolateral prefrontal cortex and anterior insula, in addition to the brain regions identified in the univariate analysis, such as the PAG and SMA, appeared to be important contributors to the prediction. In addition, different from Hoeppli et al., the Neurologic Pain Signature (NPS)^16^, an *a priori* multivariate pattern-based fMRI marker of pain, was able to explain the individual differences in pain with a significant correlation, *r* = 0.201, *p* = 0.0246 (**Fig. 2d**). Overall, our results suggest that the brain activity patterns in both univariate and multivariate analyses can capture the interindividual differences in pain ratings.

**Fig. 2.**
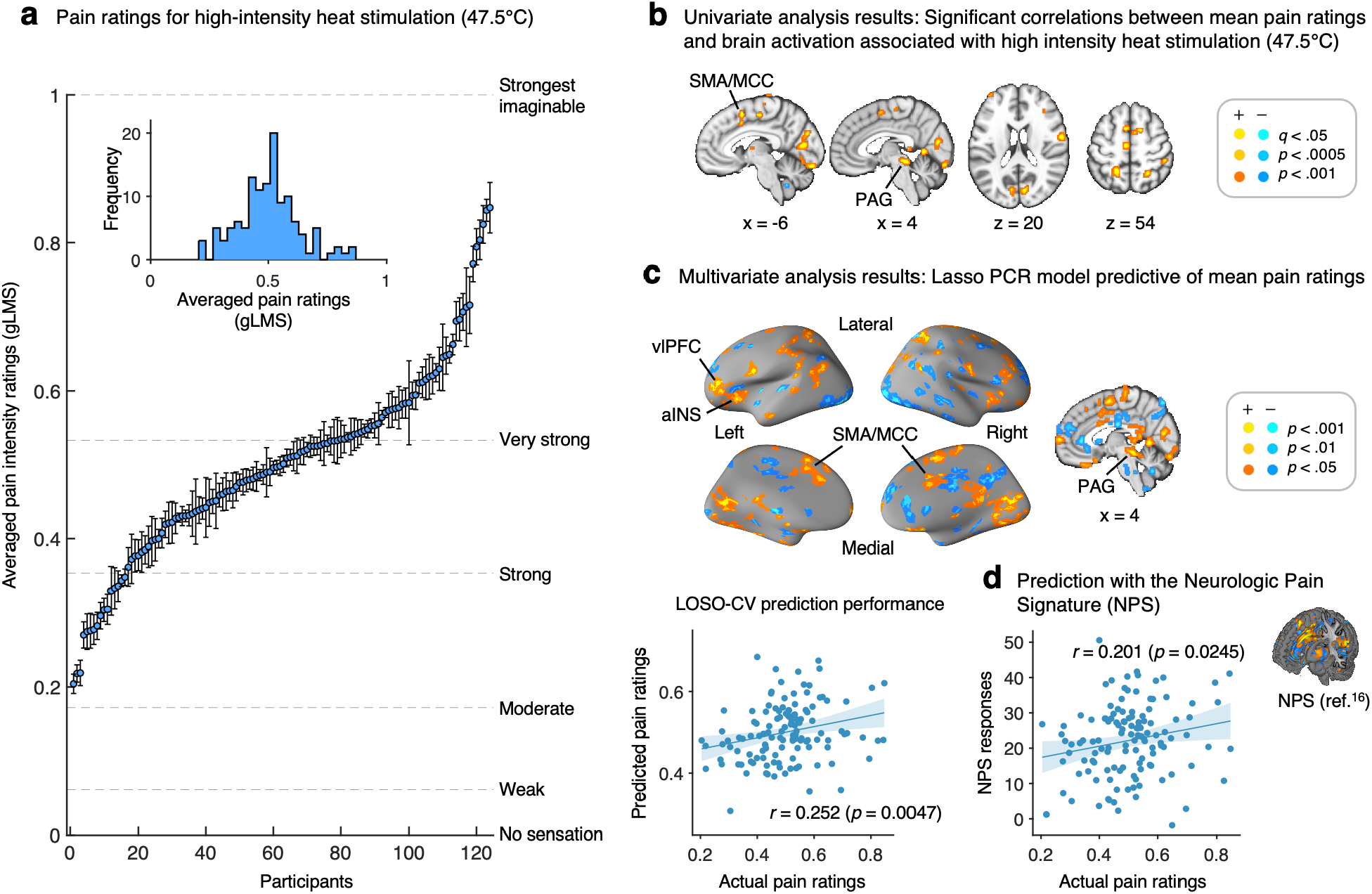
Brain activation patterns correlated with interindividual differences in pain ratings during high-intensity heat simulation. We replicated a series of analyses performed by Hoeppli et al. using a large-scale pain fMRI dataset (*N* = 124), which included a comparable number of participants to Hoeppli et al. **a**, The plot shows the distribution of average pain ratings for high-intensity heat stimulation (47.5°, the number of repeats = 16). The error bars represent the standard error of the mean for each individual’s pain ratings. The ratings were sorted in ascending order. Dashed horizontal lines indicate anchors of the general Labeled Magnitude Scale (gLMS)^15^ used in the study. **b**, Univariate analysis results with a general linear model (GLM) using individual differences in averaged pain ratings as a covariate. The input images were the brain activation maps for high-intensity heat stimulation. The brain maps were thresholded with *q* < 0.05, FDR corrected. We pruned the map using two additional more liberal thresholds, uncorrected *p* < 0.0005 and *p* < 0.001, two-tailed, to show the extent of activation clusters (see **Supplementary Fig. 1** for the main effects of high-intensity heat stimulation). **c**, Multivariate analysis results with Lasso-regularized Principal Component Regression (LASSO-PCR) to predict interindividual variations in pain ratings. (top) The map shows the voxels that reliably contributed to the prediction of mean pain ratings based on bootstrap tests (thresholded at uncorrected *p* < 0.001, two-tailed). Thresholding was performed for the purpose of display; all weights were used in the prediction. We pruned the map using two additional more liberal thresholds, uncorrected *p* < 0.01 and *p* < 0.05, two-tailed, to show the extent of clusters. (bottom) The scatter plot shows the actual versus cross-validated predicted pain ratings based on the LASSO-PCR model. The prediction-outcome correlation was 0.252, *p* = 0.0047, two-tailed. **d**, We also tested an *a priori* fMRI multivariate pattern-based marker of pain, Neurological Pain Signature (NPS^16^). Different from Hoeppli et al., which observed no relationship between the NPS response and pain ratings, the results showed a significant prediction-outcome correlation between the NPS response and the individual differences in pain ratings, *r* = 0.201, *p* = 0.0245. The scatter plot shows the actual pain ratings versus NPS responses. NPS response was calculated with the dot-product between the NPS weights and brain response to high-intensity heat stimulation. Error bands represent the 95% confidence interval of the regression lines.

There can be many reasons for the discrepancy between our results and those of Hoeppli et al., including the sample differences (e.g., sociodemographic background and psychological status) and the experimental and analysis factors, such as MR scanner, fMRI sequence, experimental design and procedure, rating scale, preprocessing steps, etc. Among these, the sample characteristics may be one of the most important factors. This idea is supported by a recent study, which reported that the brain-phenotype models reliably failed in individuals who deviated from stereotypical profiles^17^, underscoring the importance of considering sociodemographic and psychological factors to make generalizable predictive models. Hoeppli et al.’s failure to identify brain regions and models that are correlated with individual differences in pain may be due to the high heterogeneity of their sample, resulting in more complex interactions among the sociodemographic and psychological factors. In contrast, ours was more homogeneous in terms of age (mean ±SD: 22.17 ±2.69 years, range = 19-37) and educational level. While acknowledging the commendable efforts of Hoeppli et al., our findings suggest that it may be crucial to incorporate the complex interactions among the sociodemographic, psychological, and neurobiological factors (referred to as biopsychosocial factors) into analyses, particularly in cases where the sample is highly heterogeneous. Thus, although Hoeppli et al.’s null findings may initially appear discouraging, they offer deep insights into the intricate interplay among a multitude of biological, psychological, and sociodemographic factors that contribute to the interindividual variability of pain. Overall, these findings shed light on how to approach the understanding and modeling of the multifaceted nature of pain.

