## Supplementary information for "Interindividual differences in pain can be explained by fMRI, sociodemographic, and psychological factors"

**This PDF file includes:**

1. Supplementary Methods (pp. 2-4)
2. Supplementary Figure (p. 5)
3. Supplementary References (p. 6)

### SUPPLEMENTARY METHODS

#### Dataset descriptions

In this study, we utilized a large-scale dataset (*N* = 124) collected at Sungkyunkwan University. A total of 137 healthy and right-handed participants were recruited from the Suwon area in South Korea. Eligibility was assessed through online questionnaires, including pain and MRI safety-screening questions. We excluded participants with psychiatric, physiological, or pain disorders, neurological conditions, or MRI contraindications. Additionally, thirteen participants were excluded from the analysis due to technical issues with the thermal stimulus equipment, voluntary request to quit the scanning session, or the presence of abnormal brain structures (e.g., arachnoid cyst). Thus, the remaining 124 participants were included in the current dataset (female = 61, mean age = 22.2 years, SD age = 2.7 years). During the study, participants experienced contact heat stimuli and rated their pain levels after each thermal stimulation, which was delivered to the volar surface of the left forearm using a Pathways system (Medoc Ltd) with a 16-mm ATS thermode endplate. Participants rated the magnitude of their warmth or pain sensation on a general Labeled Magnitude Scale[1] following the stimulation offset. The scale was continuous ratings from 0 to 1 with anchors of "No sensation" (0), "Weak" (0.061), "Moderate" (0.172), "Strong" (0.354), "Very Strong" (0.533), and "Strongest imaginable" (1), but the anchors were not presented during the task. The temperatures ranged from 45 °C to 47.5 °C (in 0.5 °C increments, totaling 6 intensities), and the duration of each stimulation was 12 seconds (ramp-up: 2.5 secs; plateau: 7 secs; ramp-down: 2.5 secs). There were eight heat-induced task runs with twelve trials per run, resulting in 16 trials per participant for each temperature. The total number of trials was 96, but due to the errors in stimulus delivery or trials with a high variance inflection factor (> 3), the average number of trials used in the analyses was 91.6 (standard deviation = 10.6). Pre-stimulus state manipulations involved video viewing. We obtained written consent from all participants, who also received financial compensation. The same dataset was previously used in a published study[2], as an independent test dataset. The study addressed different research questions from the current study.

#### fMRI acquisition and preprocessing

The whole-brain fMRI images and high-resolution T1-weighted structural images were obtained using a 3-Tesla Siemens Prisma scanner at the Center for Neuroscience Imaging

Research, Sungkyunkwan University. We obtained functional echo-planar images (EPI) using the following sequence parameters: TR of 460 ms, TE of 27.20 ms, multiband acceleration factor of 8, field of view of 220 mm, voxel size of $2.7 \times 2.7 \times 2.7$ mm$^3$, and slice order acquisitions of interleaved. The preprocessing of the functional EPI images was performed using Statistical Parametric Mapping 12 (SPM12) and FMRIB Software Library (FSL). To ensure image intensity stability, the initial 18 volumes (approximately 8 seconds) were removed from each run. Then, the functional EPI images were corrected for motion (i.e., realignment). Distortion caused by the magnetic field inhomogeneity was also corrected using FSL's topup function. Then, the functional EPI images were co-registered and spatially normalized into the Montreal Neurological Institute normative atlas with voxel interpolation at $2 \times 2 \times 2$ mm$^3$. We then smoothed the images with a 5-mm full width at half-maximum. To reduce motion-related artifacts, we conducted an Independent Component Analysis-based strategy for Automatic Removal Of Motion Artifacts (ICA-AROMA)[3]. In addition, we excluded data from some runs based on the following two criteria regarding frame displacement (FD), which quantifies the frame-wise displacement of images: (1) the mean FD of a run exceeding 0.2 mm, and (2) the FD of any volume of a run exceeding 5 mm[4,5].

**First-level analysis: single-trial analysis**

We utilized a single-trial design approach to model the brain responses to heat stimulation. In this approach, the response magnitude of each voxel for each trial was estimated using a general linear model (GLM). This model included separate regressors for each pain trial, as in the 'beta series' approach[6]. Additional regressors, event boxcars convolved with the canonical hemodynamic response function, were created to model the periods of pre-stimulus (movie-viewing), anticipation, heat stimulation, and pain rating. Given that we already removed motion-related artifacts through ICA-AROMA during the preprocessing stage, five principal components of WM and CSF signal and a linear trend were included as nuisance covariates. Subsequently, we calculated variance inflation factors (VIFs) on a trial-by-trial basis. VIFs are a measure of design-induced uncertainty caused by collinearity with nuisance regressors. This step aimed to identify and exclude trials where the estimates could be significantly influenced by artifacts occurring during the trials. Any trials with VIFs exceeding 3 were excluded from the analyses. On average, 0.1371 trials were excluded due to high VIFs, with a standard deviation of 0.7686. Finally, single-trial beta images were obtained and served as input images for predictive modeling.

#### Univariate voxel-wise analysis with average pain intensity ratings as a covariate

We obtained the averaged beta estimates and pain intensity ratings for the highest temperature condition (47.5 °C, 16 trials) per participant. Then, we conducted a GLM analysis on the average beta images for the highest temperature and used the average pain intensity ratings as a covariate. The main effect of the highest temperature condition is shown in **Supplementary Fig. 1**, and the correlates of the average pain intensity ratings are shown in **Fig. 2b**.

#### Multivariate analysis with the Neurological Pain Signature (NPS)

We tested *a priori* fMRI multivariate pattern-based marker of pain, Neurological Pain Signature (NPS)[7], to examine whether we can predict the individual differences in pain intensity ratings with NPS responses. To obtain the NPS response, we conducted the dot-product of the NPS pattern weights and the average beta estimates for the highest temperature condition. We then calculated the correlations between the NPS response and the individual differences in the pain intensity ratings (**Fig. 2d**).

#### Multivariate analysis with LASSO-PCR

To develop an fMRI-based predictive model of the average pain intensity rating, we used lasso-regularized principal component regression (LASSO-PCR) with leave-one-subject-out cross-validation (LOSO-CV). The input features for the modeling were the average beta estimates of the highest temperature condition, and the outcome variable was the average pain intensity rating for the highest temperature stimulus. The cross-validated prediction performance of the predictive model was assessed with a prediction-outcome correlation, which refers to the correlation between the predicted and actual pain intensity ratings. To identify brain voxels reliably contributing to the prediction, we thresholded the predictive weight map using *p*-values from a bootstrap test with 5,000 iterations.

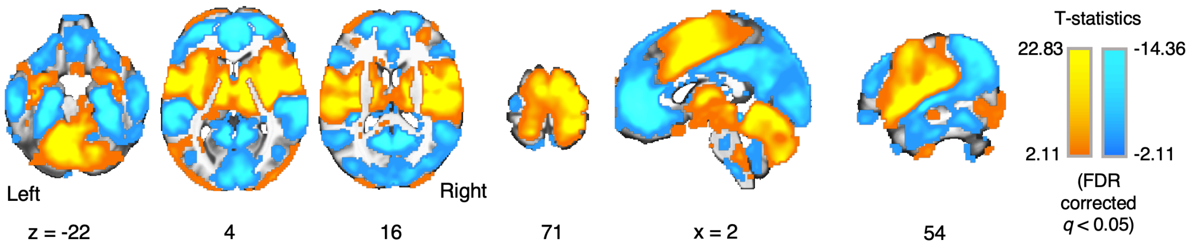


**Supplementary Figure 1. Main effect of the highest heat stimulation (47.5°C) in a univariate analysis.** The result of the univariate analysis employing a general linear model with individual differences in the average pain intensity ratings used as a covariate. The brain activation map shows the main effect of the 47.5°C heat stimulation. We applied a threshold to the map with $q < 0.05$ corrected for multiple comparisons using a false discovery rate.
